# Indistinct changes in the mammal fauna between 1986-2017 at Judbarra-Gregory National Park, Northern Territory

**DOI:** 10.1101/2025.10.09.681490

**Authors:** A.S. Kutt, D.S. Martin, A. Fisher

**Author notes:** **Target:** Austral Ecology. **Declaration of Funding:** The authors did not receive any specific funding for this project. **Data Availability Statement:** The data will be available on request.

## Abstract

The mammal fauna in many parts of northern Australia has undergone dramatic changes in the past 20-30 years, though longitudinal data that spans this period are limited and largely from coastal and high rainfall regions. We examined data from Judbarra-Gregory National Park that has been collected in a systematic manner (once in 1986 and then 2005-2017), including two years where camera trapping was included (2015 and 2017). We standardised this data in a simple manner (i.e., species by relative abundance and site frequency) to examine if there were any stark changes over this time span. We also undertook some simple modelling of mammal pattern with respect to fire and fractional green and bare cover metrics. Though we recognise some limitations in our comparisons (i.e., spatial variation in sites and methods), the data indicated that the mammal fauna had generally not changed substantially in composition, relative abundance or frequency across sites. There were meaningful models correlating mammal pattern to fire frequency, time since fire, fraction green and bare parameters. However, some species symptomatic of small mammal decline elsewhere (i.e., *Rattus tunneyi*) may have become less common, whereas other species were recorded for the first time in more recent years (2010-2017), particularly by camera traps (i.e., *Trichosurus vulpecula, Isoodon macrourus, Melomys burtoni*). We postulate that change in the mammal fauna on the semi-arid fringe of tropical savannas may have occurred prior to the 1980s, with a long history of cattle grazing a possible contributing factor. The mammal fauna there, is also perhaps naturally sparse and heterogeneous in spatial and temporal distribution.

## INTRODUCTION

The mammal fauna of some parts of the Top End of the Northern Territory has undergone a calamitous decline in the past three decades, Kakadu National Park providing the most well documented example of this pattern (Woinarski *et al*. 2010). Though there is evidence in other regions of northern Australia of an equivalent contemporary depletion of mammals (McKenzie *et al*. 2024), in many cases detailed temporal data is absent (Kutt *et al*. 2012). The causes for the decline are sometimes readily articulated (i.e., feral cat predation, cane toad poisoning, and habitat loss and decline in habitat condition due to inappropriate fire regimes, Woinarski *et al*. 2015), especially when the timing of decline is clear from long term monitoring. For other examples of changing abundance of species, the causes are more conceptual and uncertain (Radford *et al*. 2020).

In the more southern portions of the tropical savannas, there is a prospect that changes in the contemporary mammal fauna commenced earlier than the well reported declines in the northern near-coastal savannas, that occurred the start of this century (Woinarski *et al*. 2010). Periods of substantial drought coupled with extensive grazing by introduced herbivores may have contributed to an earlier phase of loss. The absence of long-term data to demonstrate changes in mammal populations across this period (i.e., 1950’s to the present) means that in many parts of northern Australia, the reconstruction of the changes in fauna and their causes is correlative guesswork (Kutt and Fisher 2011; Kutt *et al*. 2012; Woinarski *et al*. 2006). There are simple descriptive accounts that indicate that older fauna inventories recorded very few small mammals (i.e., unpublished data from surveys described in Kirkpatrick and Lavery 1979) and that in Queensland, at least, there were extended periods of substantial pasture degradation (McKeon *et al*. 2004). Of course, without associated monitoring data, collected in concert with potential drivers of mammal change, the causes of any transformation noted is in part speculative.

In this review of previously unpublished data spanning the period 1986 to 2017, we investigate whether there are any apparent changes in the mammal fauna in a very large conservation reserve that spans the sub-tropical to semi-arid zone in the western Northern Territory. We examined simple metrics of mammal abundance and frequency to examine if any change occurred over this 31-year period and examine if there is any modelled correlation to fire patterns and vegetation cover. The data we examined was not collected for, or designed to, investigate environmental correlates of change, so we treat our analyses cautiously. Understanding the variation in patterns of mammal abundance across a range of geographic and climatic locations in northern Australia, and over time, may assist in better disentangling the temporal and spatial extent of contemporary mammal declines, and to better inform applied conservation management.

## METHODS

The data were collected in Judbarra-Gregory National Park (JGNP, centroid: -16°24’ S, 130° 24’ E) in the Northern Territory, 360 km south of Darwin. The park is the second largest national park in the Northern Territory (1 299 455 hectares), after Kakadu National Park, and was gazetted in August 1990. It spans an area from Timber Creek in the north to Kalkarindji in the south, a range in mean annual rainfall from 970 to 690mm. The vegetation is predominantly open woodlands (tropical savanna to semi-arid woodlands) but includes woodlands and heath on dissected rocky plateaus and escarpments, monsoon vine-thicket, riparian forest and grasslands.

We examined data from nine fauna surveys conducted at JGNP from 1986 to 2017, that used methods summarised in Supplementary Table 1. Further descriptions of each method can be found in Einoder *et al*. (2020) and (Woinarski *et al*. 2010). We only considered mammal data, although reptiles, amphibians and birds were surveyed as well. The methods and trapping effort differed slightly between surveys, but we tabulated the mean number of nights each site was surveyed (and maximum and minimum), and for each technique the total traps per sites, or hours searched (with the maximum and minimum) (Supplementary Table 1). For our analyses, we assumed that the effort was comparable from year to year, recognising the greatest variation was in the first survey where the Elliott trapping effort was higher, and no diurnal active searching occurred. Therefore, for each mammal recorded we simply used the mean abundance across all sites (and standard error) and, as a more conservative metric, the frequency of occurrence for each species across all the sites surveyed.

Camera trapping only occurred in 2015 and 2017, and we present this data separately from the conventional trapping and search data, for comparison. Scientific and common names follow the Northern Territory Fauna Species Checklist (https://data.nt.gov.au/dataset/nt-fauna-species-checklist).

We modelled the species richness (native and introduced) and abundance of all native and introduced mammals, and mammal species (recorded in at least 5 sites in total) separately using generalised linear mixed models (GLMMs) with a negative binomial error distribution to account for overdispersion and the high frequency of zeros (Brooks et al. 2017). Four habitat and disturbance predictors were considered: fractional cover of green vegetation (FC_green), fractional cover of bare ground (FC_bare), fire frequency (FireFq), and time since last burn (TSLB). Fire data were produced by North Australia and Rangelands Fire Information (NAFI) at Charles Darwin University, derived from satellite imagery sourced from the Moderate Resolution Imaging Spectroradiometer (MODIS) using the Red and Near Infrared bands with a 250 × 250 m pixel resolution. The fire metrics were calculated using standard GIS techniques covering the 10 years prior to each of the survey years. The fire metrics were attributed to each site using the mean value of a 3 × 3-pixel window centred over the survey location.

The fractional cover data is the estimated proportion of bare ground (i.e., soil) and green (i.e., photosynthetic vegetation in the ground, shrub, and tree layer), within each pixel. For this study, we used the three-month seasonal Landsat composite product, which selects representative pixels through the determination of the medoid (multi-dimensional equivalent of the median) of three months (a season) of fractional cover imagery (Joint Remote Sensing Research Program 2021). For further details on this compositing method see Flood (2013). The product was extracted via the VegMachine API (Beutel *et al*. 2019), which now uses an updated version of fractional cover products (Joint Remote Sensing Research Program and Department of Environment and Science 2022).

All continuous predictors were standardised (mean = 0, SD = 1) prior to analysis to facilitate comparison of effect sizes. Survey year was included as a random intercept to account for temporal non-independence among repeated sampling occasions. We used an information-theoretic approach to identify the most parsimonious predictor set, fitting all possible subsets of the four predictors (16 candidate models per species) and ranking models by corrected Akaike’s Information Criterion (AICc) (Burnham and Anderson 2002). Model-averaged coefficients and unconditional standard errors were derived from the confidence set of models (ΔAICc ≤ 4) using full model averaging, whereby predictors absent from a given model were assigned a coefficient of zero. The relative importance of each predictor was calculated as the sum of AICc weights across all models in which it appeared. Residual diagnostics were performed using the DHARMa package to assess model fit, overdispersion, and zero-inflation (Hartig 2016). Where substantial zero-inflation was detected, models were refit with an additional zero-inflation component. All analyses were conducted in R 4.5.2 (R Core Team 2024) using the packages glmmTMB (Brooks *et al*. 2017) and MuMIn (Bartoń 2024).

## RESULTS

The simple analysis of mammal data recorded at Judbarra-Gregory National Park using broadly consistent methods from 1986 to 2017, found limited variation in species richness, mean abundance and frequency across this time span. There appeared to be some depletion of mammals after 1986, though improvement in 2012-2017 (Table 1). Camera trapping resulted in equally high species richness in 2017 compared to 1986. The mean abundance of all mammals showed little notable change, though with perhaps some reduction in the period 2005-2011; however, the total and mean number of introduced mammals increased over time. The frequency of native mammal captures did not vary in any meaningful way over time (Fig. 1).

**Table 1.**
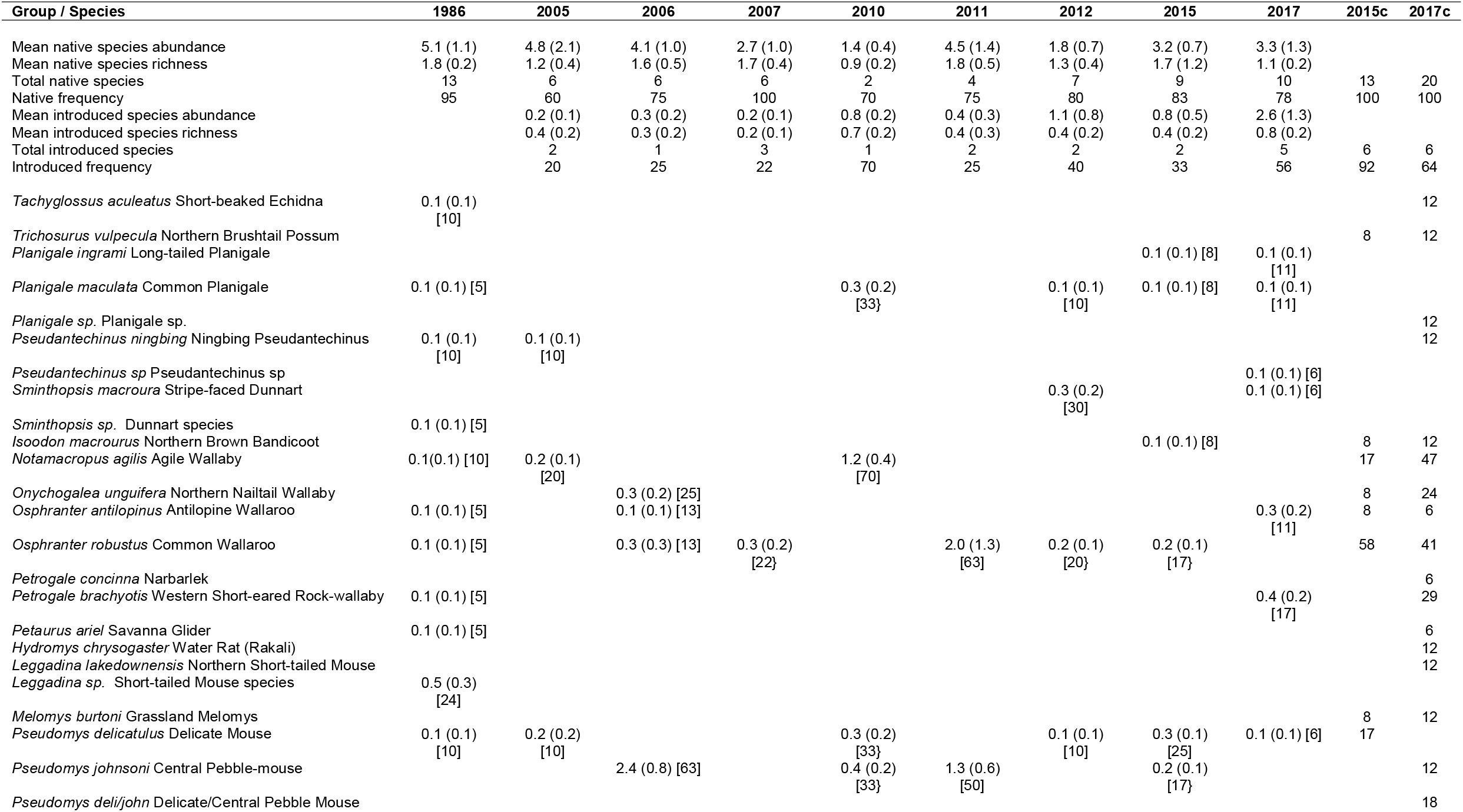
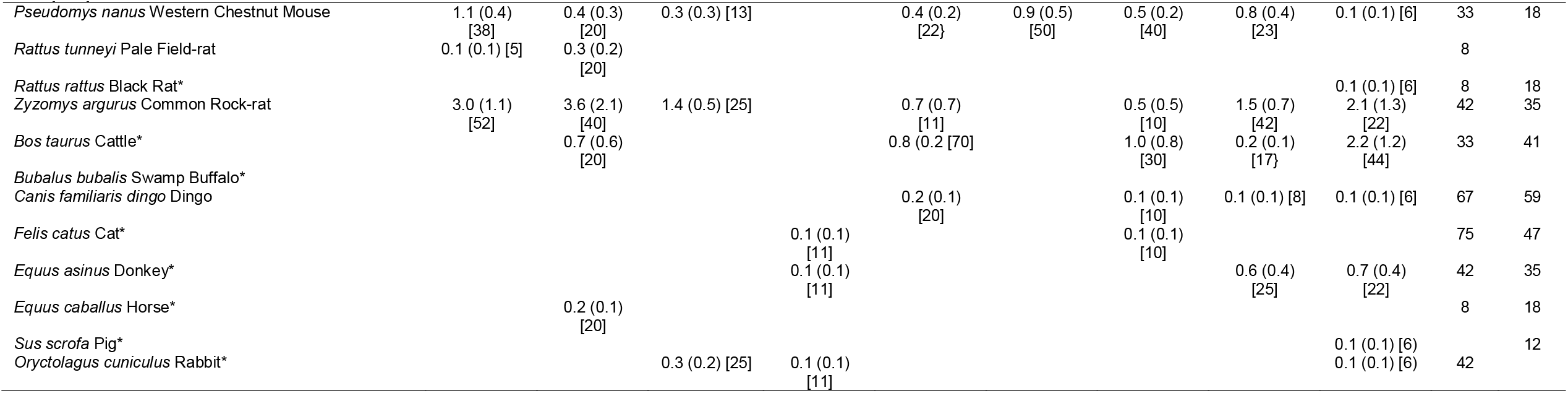
The mean (and S.E) of abundance and species richness of all native and introduced mammals per site for each year, and the total number of native and introduced species for each survey. Native frequency and Introduced frequency are the proportion of sites for that year from which they were recorded. For the individual species, the data is the mean (and S.E.) site abundance and the frequency across sites (square brackets). The camera data (2015c and 2017c) is the total native and introduced species recorded, the frequency of these two groups and each species, recorded across the camera traps set.

**Figure. 1.**
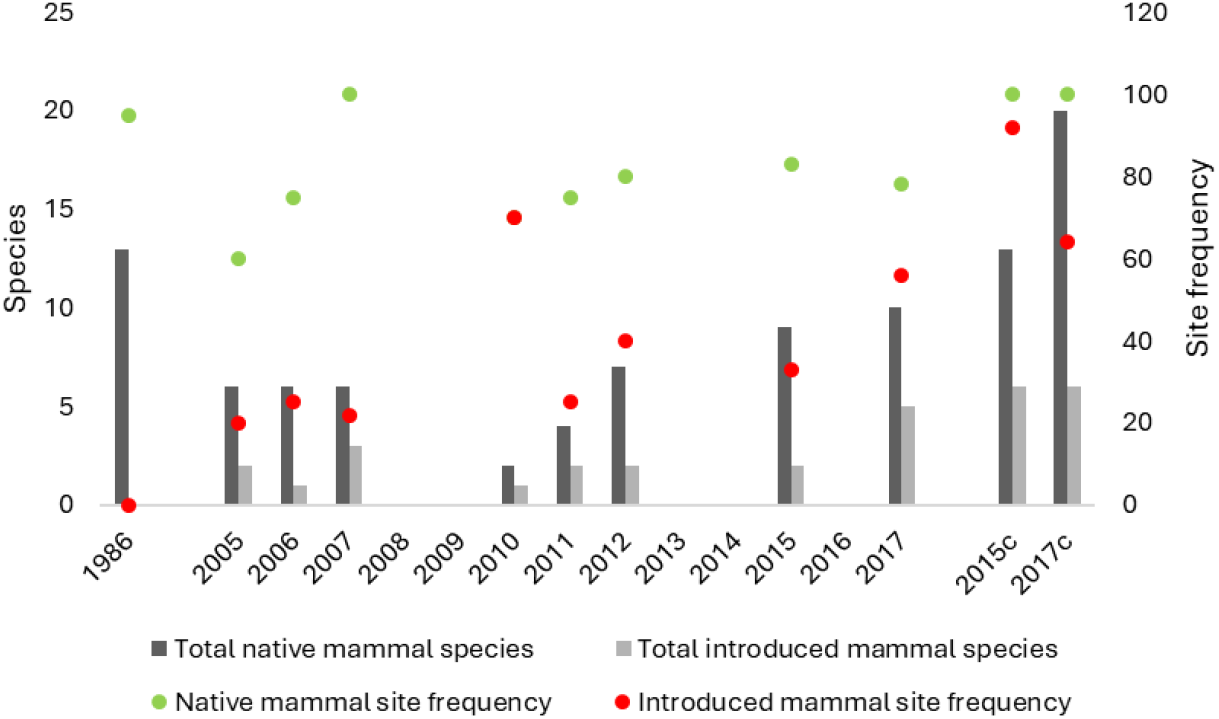
The frequency of sites in which native and introduced mammals were recorded in each survey (represented by the green and red markers), and the total species richness of native and introduced mammals recorded for each survey (represented by the bars).

We also examined the pattern in four environmental variables (two fire metrics and two fractional cover metrics that coarsely reflect vegetation structure and response to rainfall or herbivory) (Fig. 2 a, b). In the case of the fractional cover green and bare ground, there was not a great deal of variation in the average for each year for the survey sites (between 40-50% green cover, 15-30% for bare ground). Similarly, the fire frequency and time since last burnt was relatively consistent, except for a lower fire frequency in 2006 and 2007. Model-averaged coefficients indicated weak and uncertain associations between all four habitat predictors and mammal groups or species, with 95% confidence intervals overlapping zero in every case where there was sufficient data to model (native and introduced mammal abundance, and four species, Supplementary Table 2). Relative importance weights were consistently low across all predictors and species indicating that no single predictor was strongly or consistently supported across the candidate model set. Six mammal species and groups were precluded from reliable model fitting and parameter estimation due to insufficient data or lack of variation in abundance across the range of environmental variables sampled (i.e., low number of sample sites in each year. This suggest that at least for mammal data collected at JGNP, it is not strongly structured by fire or vegetation cover, and that substantial year to year variation in sampling, may be obscuring any environmental associations.

**Figure. 2.**
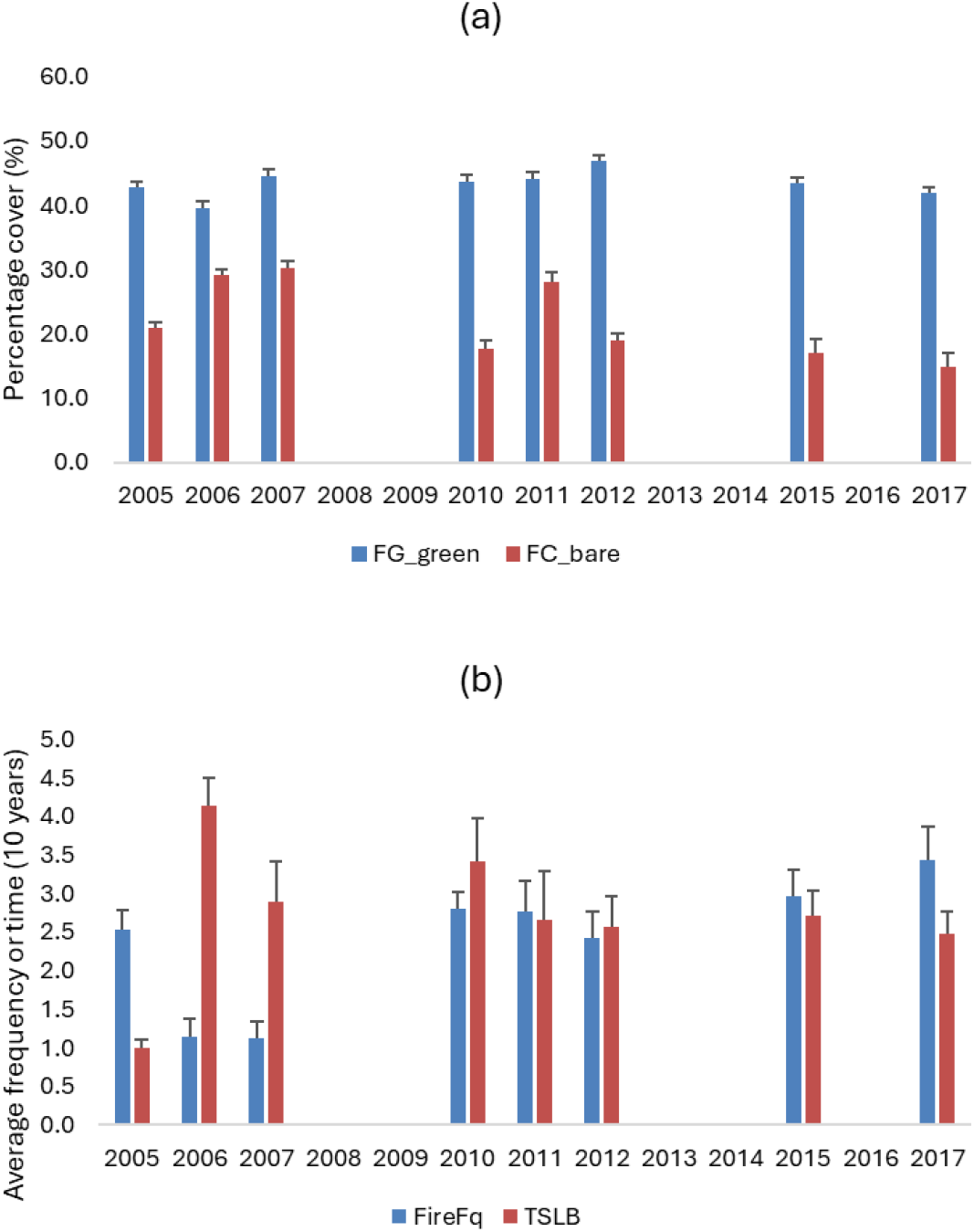
The mean (and S.E) of (a) fractional cover green (FC_green) and fractional cover bare (FC_bare), and (b) fire frequency and time since last burn for each survey year (2005-2017). These data were unavailable for 1986.

## DISCUSSION

The intent of this short paper was to investigate any distinctive changes in the mammal fauna (species or abundance) at Judburra-Gregory National Park, over multiple years of survey. The intent of the original data collection was to inventory the fauna at the park, and not to investigate management such as fire regimes, though this has occurred for vegetation (Russell-Smith *et al*. 2010). We included modelling of mammal pattern in response to fire and fractional cover metrics at the request of anonymous reviewer, even though this seemed like overreach for a data set more suitable for more gentle examination. The lack of any meaningful results reaffirmed the difficulty in retrofitting data collected for one purpose (i.e., surveillance, discovery or pattern analysis) to landscape scale management questions (Kutt *et al*. 2026). At a simple level, the data indicated that in some geographic locations (i.e., semi-arid savanna), mammal composition and abundance has not changed markedly over the past two decades, older outliers (i.e., 1986 data) point to similarly low abundance of mammals, and novel methods such as camera trapping, provide better options to systematically understand contemporary mammal pattern.

We recognise a number of other limitations in our study and thus the interpretation of the data, namely; (i) the slight variation in methods and effort over the years, though given the low number of mammal captures, we believe this is inconsequential, (ii) the lack of repeat sampling at sites, which changed annually, and therefore might have sampled different vegetation types, and different locations across the very large JGNP. This could influence temporal trends recorded across a large, heterogeneous park (though there were not meaningful trends), and (iii) the variation in weather patterns and survey timing from year to year (though all sampling occurred in the dry season between May and August). However, given the unambiguous changes in Kakadu from 1986-2010 (Woinarski *et al*. 2010), and the straightforward analysis of that data (i.e., due to the magnitude of change) we expected to see a similar dramatic change evident from the 1986 data, if an equivalent mammal decline was occurring. What is most notable from the data is the generally low abundance and frequencies of all mammal species recorded, even in 1986 when the trapping effort was very high.

Smaller native species (< 5.5 kg) are the cohort considered to be reducing in abundance and range in tropical savannas (Fisher *et al*. 2014). In examining the pattern in smaller native species in our data set, the abundance of species such as *Pseudomys delicatulus, P. nanus*, and *Zyzomys argurus* did not seem to change over the years of survey. Other species such as *Leggadina* sp. can be naturally variable in occurrence (Kutt and Kemp 2005). One species did seem to show a typical pattern of decline (McKenzie *et al*. 2024); *Pseudomys tunneyi* was not recorded after 2005 by conventional trapping, and when recorded by camera trapping in 2015 it occurred in very few sites. Black rats *Rattus rattus* appeared for the first time in 2017, and may affect the persistence of native mammals (Banks and Hughes 2012). Curiously, a handful of species that are commonly recorded in northern Australia were only documented for the first time on camera traps, or in surveys in more recent years (Stripe-faced Dunnart *Sminthopsis macroura*, Northern Brushtail Possum *Trichosurus vulpecula*, Northern Brown Bandicoot *Isoodon macrourus*, Grassland Melomys *Melomys burtoni*, Rakali *Hydromys chrysogaster*). This may partly reflect the limitation that different sites were surveyed each year and habitat specialists (*H. chrysogaster, Pseudomys johnsoni, Pseudantechinus* sp.) may be unrecorded due to sampling of unsuitable habitat in any one year. Camera trapping in general allows a better inventory of more cryptic species (Thomas *et al*. 2020), though it is best used in concert with conventional trapping and manual search methods (Kutt *et al*. 2023).

This review of survey data, although in part descriptive, demonstrated limited change in the GJNP mammal fauna across the span of the surveys, with many species only infrequently reported. Introduced species seemingly increased over the survey years, (i.e., Feral Cat *Felis catus*, Black Rat *Rattus rattus*, and Feral Pig *Sus scrofa*) though these species are recorded inefficiently using the trap and search methods described in this study; this pattern may not reflect any change except for Black Rats which are now recorded with increasing abundance in tropical savannas. Although there are limitations on the interpretation of this data, a dramatic decline in the mammal fauna comparable to some other well-studied Top End locations was not evident, and we hypothesise that, due to the long history of grazing in large pastoral properties in the sub-monsoonal and semi-arid areas of the tropical savannas, the most definitive changes in the mammal fauna may have already occurred prior to the 1980s (Kutt and Fisher 2011). In addition, our attempt to model mammal pattern with typical predictors of their occurrence or change (i.e., fire, vegetation cover) highlighted the spareness of the JGNP data; data collected for a different purpose. As such, more carefully stratified surveys and landscape scale ecological experiments to examine the cause and timing of contemporary declines and the ongoing trajectory of mammals in geographically contrasting locations should continue to occur as a priority in northern Australia.

## ACKNOWLEDGEMENTS

The data used in this study have been derived from a wide range of field surveys and research programs funded by the Northern Territory and Australian Governments. Many staff from the Flora and Fauna Division, NT Parks and Wildlife and other volunteers assisted with the data collection over the span of the surveys.

---

**Supplementary Table 1.** The number of sites and methods used for each of the surveys examined. The data is the total per site and in parentheses, the maximum and minimum (or the total reiterated if numbers didn’t vary across the sites). Diurnal and nocturnal are active searches, and the data is person-hours. Cameras (total) are the number set per site and the total for the survey in parenthesis. Camera nights are the mean number of nights the cameras were set, and the maximum and minimum in parenthesis.

| Year | Sites | Nights | Elliott | Cage | Pitfall | Diurnal | Nocturnal | Cameras<br>(total) | Camera<br>nights |
| --- | --- | --- | --- | --- | --- | --- | --- | --- | --- |
| 1986 | 21 | 3.6 (3-4) | 100.0 (100) | 4.0 (0-20) | 1.4 (0-10) | 0 | 0.6 (0-2) | 0 | 0 |
| 2005 | 10 | 3.0 (3) | 20.0 (20) | 4.0 (4) | 3.8 (3-4) | 3 (3) | 2 (2) | 0 | 0 |
| 2006 | 8 | 3.0 (3) | 20.0 (20) | 4.0 (4) | 4 (4) | 3 (3) | 2 (2) | 0 | 0 |
| 2007 | 9 | 3.0 (3) | 20.0 (20) | 4.0 (4) | 4 (4) | 3 (3) | 2 (2) | 0 | 0 |
| 2010 | 10 | 3.0 (3) | 20.0 (20) | 4.0 (4) | 4 (4) | 3 (3) | 2 (2) | 0 | 0 |
| 2011 | 8 | 3.0 (3) | 20.0 (20) | 4.0 (4) | 4 (4) | 3 (3) | 2 (2) | 0 | 0 |
| 2012 | 10 | 3.0 (3) | 20.0 (20) | 4.0 (4) | 4 (4) | 3 (3) | 2 (2) | 0 | 0 |
| 2015 | 12 | 4.0 (4) | 16.0 (16) | 8.0 (8) | 3.4 (2-4) | 3 (3) | 2 (2) | 7 (84) | 82.1 (80-85) |
| 2017 | 18 | 3.9 (1.5-4) | 16.0 (16) | 8.0 (8) | 3 (3) | 2.7 (0-3) | 2.5 (2-4) | 5 (90) | 49.1 (42-55) |

**Supplementary Table 2.** The six mammal groups or species where reliable model fitting and parameter estimation could occur. FC_green is fractional cover green, FC_bare is fractional cover bare, TSLB is time since last burn and FireFq is fire frequency Z(the latter two, being measured in the last 10 years prior).

| Groups / species | Predictor | Estimate | SE | Lower 95%<br>CI | Upper 95%<br>CI | Importance | z | p |
| --- | --- | --- | --- | --- | --- | --- | --- | --- |
| Native mammal abundance | FC_green | 0.008 | 0.056 | -0.101 | 0.117 | 0.26 | 0.141 | 0.888 |
| Native mammal abundance | FC_bare | -0.004 | 0.048 | -0.098 | 0.091 | 0.25 | 0.073 | 0.942 |
| Native mammal abundance | TSLB | 0.003 | 0.049 | -0.092 | 0.098 | 0.25 | 0.053 | 0.958 |
| Native mammal abundance | FireFq | 0 | 0.047 | -0.092 | 0.091 | 0.25 | 0.008 | 0.994 |
| Introduced mammal abundance | FC_bare | 0.341 | 0.4 | -0.443 | 1.124 | 0.56 | 0.847 | 0.397 |
| Introduced mammal abundance | FireFq | 0.228 | 0.348 | -0.453 | 0.910 | 0.47 | 0.652 | 0.514 |
| Introduced mammal abundance | TSLB | 0.062 | 0.227 | -0.383 | 0.508 | 0.29 | 0.272 | 0.786 |
| Introduced mammal abundance | FC_green | -0.006 | 0.128 | -0.256 | 0.245 | 0.25 | 0.045 | 0.964 |
| Macropus robustus | FC_green | 0.421 | 0.506 | -0.569 | 1.412 | 0.56 | 0.828 | 0.408 |
| Macropus robustus | FC_bare | 0.319 | 0.499 | -0.660 | 1.298 | 0.44 | 0.635 | 0.526 |
| Macropus robustus | TSLB | 0.172 | 0.338 | -0.490 | 0.835 | 0.38 | 0.506 | 0.613 |
| Macropus robustus | FireFq | -0.016 | 0.288 | -0.580 | 0.548 | 0.27 | 0.056 | 0.955 |
| Pseudomys johnsoni | FC_bare | -0.173 | 0.569 | -1.287 | 0.942 | 0.30 | 0.301 | 0.764 |
| Pseudomys johnsoni | FireFq | 0.092 | 0.295 | -0.487 | 0.670 | 0.29 | 0.307 | 0.759 |
| Pseudomys johnsoni | TSLB | -0.096 | 0.282 | -0.649 | 0.456 | 0.30 | 0.338 | 0.735 |
| Pseudomys johnsoni | FC_green | 0.063 | 0.254 | -0.434 | 0.560 | 0.28 | 0.245 | 0.807 |
| Pseudomys nanus | FireFq | 0.153 | 0.265 | -0.366 | 0.673 | 0.43 | 0.573 | 0.566 |
| Pseudomys nanus | TSLB | -0.038 | 0.169 | -0.369 | 0.292 | 0.28 | 0.224 | 0.823 |
| Pseudomys nanus | FC_green | 0.028 | 0.122 | -0.212 | 0.267 | 0.28 | 0.223 | 0.824 |
| Pseudomys nanus | FC_bare | 0 | 0.136 | -0.268 | 0.267 | 0.25 | 0.003 | 0.997 |
| Sminthopsis macrurous | TSLB | 0.228 | 0.46 | -0.674 | 1.129 | 0.38 | 0.491 | 0.623 |
| Sminthopsis macrurous | FC_bare | -0.422 | 0.904 | -2.194 | 1.349 | 0.40 | 0.463 | 0.644 |
| Sminthopsis macrurous | FC_green | 0.186 | 0.369 | -0.538 | 0.910 | 0.38 | 0.500 | 0.617 |
| Sminthopsis macrurous | FireFq | -0.174 | 0.521 | -1.195 | 0.847 | 0.31 | 0.330 | 0.741 |

